# Protein interaction probability landscapes for yeast replicative aging

**DOI:** 10.1101/2020.08.19.257279

**Authors:** Hao-Bo Guo, Mehran Ghafari, Weiwei Dang, Hong Qin

**Affiliations:** Department of Computer Science and Engineering; SimCenter; Department of Biology, Geology and Environmental Science The University of Tennessee at Chattanooga, Chattanooga, Tennessee 37405, USA; Huffington Center on Aging, Baylor College of Medicine, Houston, Texas 77030, USA

**Author notes:** Correspondence (HG) and (HQ).

## Abstract

We proposed a novel probability landscape approach to map the systems-level profile changes of gene networks during replicative aging in *Saccharomyces cerevisiae*. This approach enabled us to apply quasi-potentials, the negative logarithm of the probabilities, to calibrate the elevation of the landscapes with young cells as a reference state. Our approach detected opposite landscape changes based on protein abundances from transcript levels, especially for intra-essential gene interactions. We showed that essential proteins play different roles from hub proteins on the age-dependent landscapes. We verified that hub proteins tend to avoid other hub proteins, but essential proteins are attractive to other essential proteins. Overall, we showed that the probability landscape is promising for inferring network profile change during aging and that the essential hub proteins may play an important role in the uncoupling between protein and transcript levels during replicative aging.

## Introduction

Studies of replicative lifespan (RLS) in budding yeast, *Saccharomyces cerevisiae*, provided mechanistic insights on cellular aging^1^, and contributed to understanding of aging in other organisms including humans^2–5^. Deletions of single genes, such as *FOB1*^6^, and suppression or overexpression of genes, such as *SIR2*^7,8^, could change the yeast RLS. Other factors that affect lifespan, such as caloric restriction (CR), were also found dependent on the genotypes^9^. A recent work found overexpression of both *SIR2* and *HAP4* genes leads to a cell state with considerably long RLS^10^.

Transcriptional fidelity is associated with aging and longevity in yeast^11,12^. The translational efficiencies of different proteins vary up to hundreds of folds in yeast, however, protein and mRNA levels were highly correlated^13^. Increased noises in both transcriptome and proteome during aging could be part of the cascading events leading to replicative aging and eventual cell death.

Protein functions are often mediated by physical interactions, especially for proteins in similar functional modules^14^. These protein interactions in the cell collectively constitute the protein-protein interaction network (PIN), a graph composed of proteins as vertices and interactions as edges^15–17^. The protein-protein interactions (PPIs) are important factors in the genotype-to-phenotype relationships^18^. The aging phenotype also intertwines with the PIN: The topology of the PIN will be altered by aging along with changes in protein abundances and/or localizations. Based on the age-dependent gene expressions in both fruit-fly and human brain, it was proposed that aging preferentially attacks key regulatory nodes important for the stability of the PIN^19^. Age-specific (or “dynamic”) PIN was found to suggest key players in aging^20^.

It was shown that cellular aging is an emergent phenomenon arising from a PIN with stochastic loss of interactions to essential proteins^21^. Intuitively, along aging, the stochastic loss of interactions associated with an essential protein may be equivalent to the deletion of the essential protein that will lead to the cell death. A recent study showed that the uncoupling between protein and transcript levels is strongly connected to yeast aging^22^. The time-series transcriptome and proteome data of this work also involved extensive numbers of essential proteins and genes^22^.

Here, using the age-dependent transcriptome and proteome dataset^22^ and a resent update of the yeast PIN^23^, we constructed the Protein Interaction Probability Landscapes (PIPLs). This method allowed us to reveal opposite landscape changes from protein levels and those based on transcript levels and suggest that interactions to essential hub proteins may play an important role in the uncoupling of proteome and transcriptome during aging.

## Result

We hypothesize that the uncoupling of proteome and transcriptome in replicative aging may lead to protein-protein interaction profile changes, which motivated us to construct age-dependent protein interaction probability landscapes (PIPLs) using the age-dependent proteome and transcriptome data along the yeast replicative aging^22^.

### Interaction potential landscape on lifespans suggest roles of essential genes for uncoupling of protein and transcript levels during aging

To gauge changes of the interaction profiles during replicative aging, we constructed the age-dependent PIPLs on dimension of replicative lifespan (RLS). We used RLS ratios between the RLS of single-gene deletion mutants and those of the wild type cells cultivated under the same YPD (2% glucose) conditions, which is referred to as the normalized RLS in present work. The normalized RLS of essential proteins (genes) are 0, as the cell will not be viable after deletion of a single essential gene. The probabilistic nature of the landscape enabled us to apply quasi-potentials to quantify the elevations of the landscape. We choose the young cells collected at the age of 7.8 hr as the ground state (Figure 1), a term borrowed from the embryonic stem cell studies^24^. Note that the quasi-potential used for the landscapes was calculated as the negative logarithm of the interaction probability based on the law of mass action (*Methods and Materials*), which bears the Helmholtz free energy (i.e., potential of mean force) characteristics^25^. Therefore, as compared with the ground state, the ridges on the landscape indicate that the interacting pairs need to acquire “high potential”, whereas basins represent that the interacting pairs possess “low potential” for interactions to occur.

**Figure 1.**
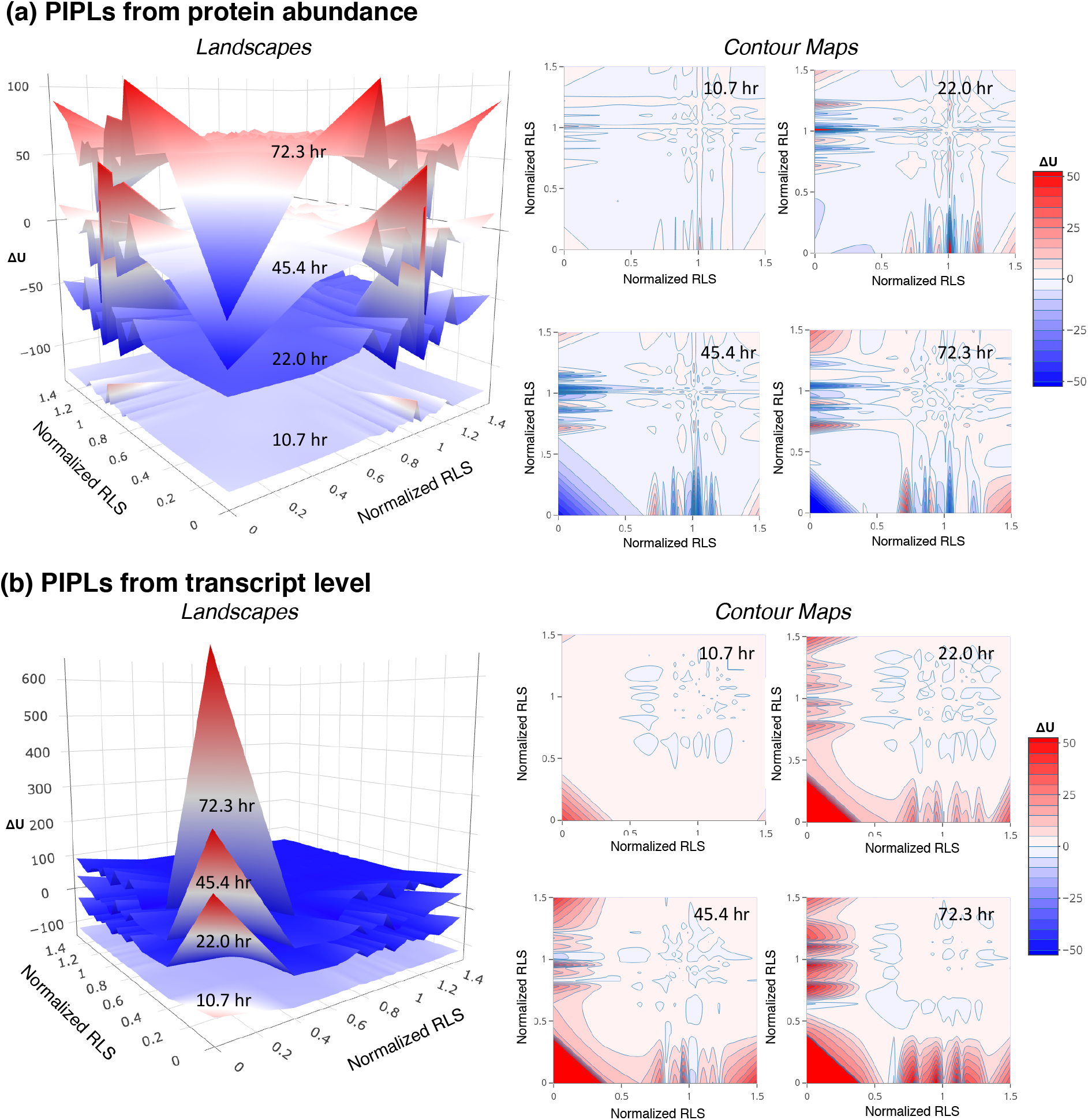
Opposite trends of intra-essential protein interactions during replicative aging as shown by the age-dependent protein interaction probability landscapes (PIPLs) on dimension of normalized RLSs. (**a**) Elevations estimated by proteome on landscapes (left) and corresponding contours (right). (**b**) Elevations estimated by transcriptome on landscapes (left) and corresponding contours (right). The relative quasi potentials (ΔU) were calculated using the youngest cells at 7.8 hr as the reference ground state. The horizontal axes of the landscapes are the RLSs of the single-gene mutants normalized by the WT measurement at the same conditions (normalized RLSs). The essential genes have normalized RLSs of 0. The landscape based on protein abundance shows that the essential-vs-essential interaction region gradually fell into a low-potential basin. However, in the landscape based on gene expression levels the essential-vs-essential interaction region gradually rose up to a high-potential ridge. The landscapes at different ages have been vertically shifted for a better visualization. The color bar shows the relative quasi potentials in the contour maps. A relative color scheme (blue for low and red for high) is applied in landscapes.

To estimate the changing elevations on landscapes during aging, we used two approaches: One based on the proteome (Figure 1a) and one based on the transcriptome (Figure 1b). We observed some strikingly opposite trends in the two landscapes for intra-essential interactions, especially at old ages. On the PIPL estimated from proteome, we found that intra-essential interactions form basins in old ages, corresponding to higher interaction probabilities than the ground state young cells. On the PIPL estimated from transcriptome, we found that intra-essential interactions form peaks in old ages. The PIPLs at selected age had been plotted, and we also visualized this opposite trend in corresponding two-dimensional contour maps (Figure 1).

### Interaction potential landscape examined by node degrees and normalized RLS

To better understand the role of essential genes during aging, we examined the interaction potential landscapes by node degrees (Figure 2a), because it was suggested that essential genes were often highly connected hubs in the protein networks^26^. Elevations of the landscape were estimated again with both protein and transcript levels. Based on protein abundances, in the old cells at 72.3 hr, the hub-versus-hub protein interactions also fell into a basin (ΔU = –31) with the quasi-potential weaker than the essential-versus-essential protein interactions (ΔU = –83) shown in Figure 1. The relative quasi-potentials for younger cells were even positive for younger cells from 10.7 to 22 hr (ΔU = 0.5 to 1.8). The extents of the quasi-potentials at other regions were also smaller than those in the PIPL based on normalized RLSs in Figure 1. For the PIPL based on transcript levels, the interactions with hub proteins, both from other hub and non-hub proteins, generally rose to ridges of the landscape.

**Figure 2.**
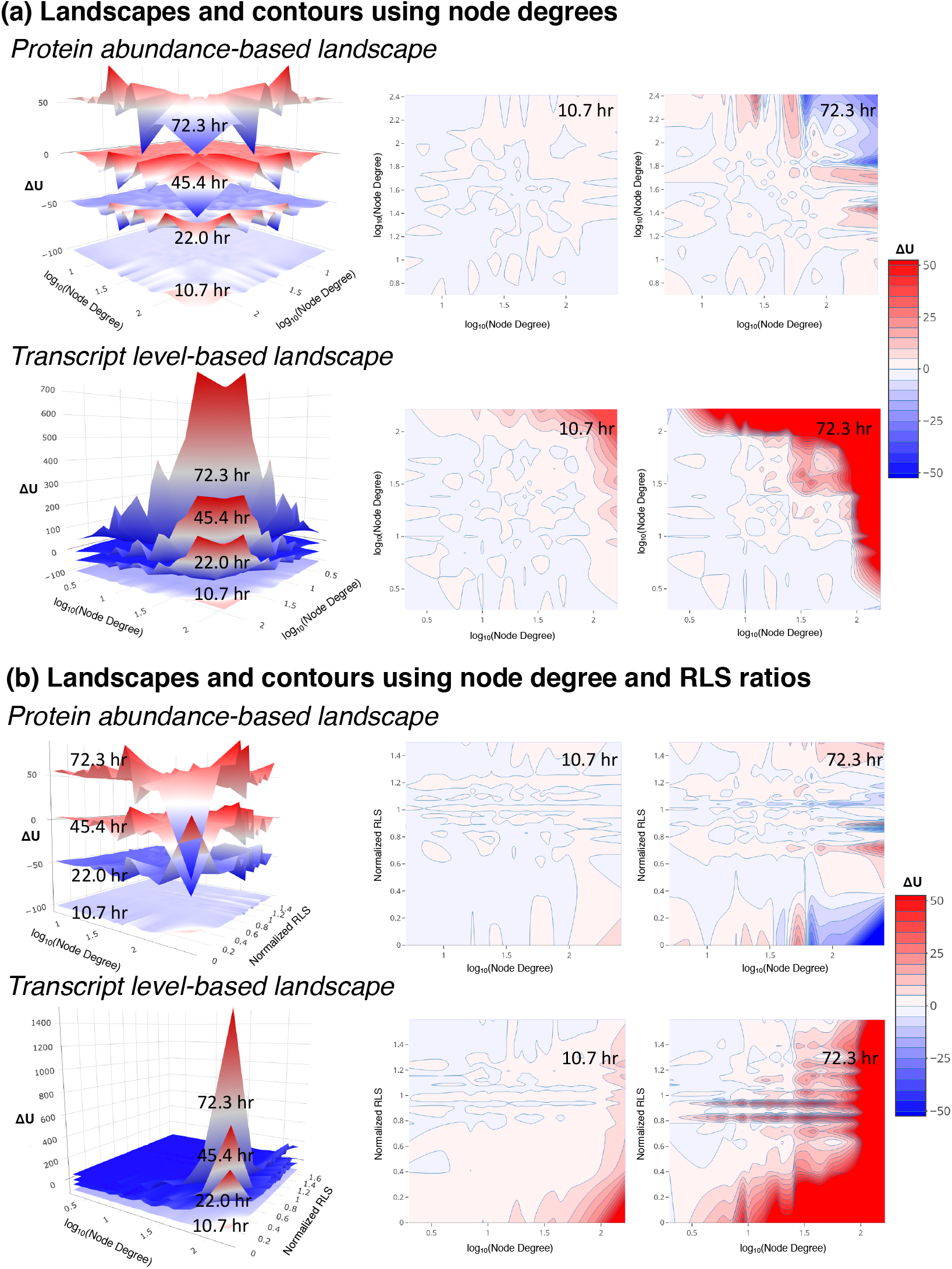
Dissimilarity between the essential and hub proteins from the PIPLs on dimensions of node degrees and normalized RLSs. (**a**) PIPLs on node degrees, and (**b**) PIPLs on node degrees versus normalized RLSs. The top panels show the PIPLs from protein abundances, and the bottom panels show the PIPLs from transcript levels. The contour maps of the aging cells at young (10.7 hr) and old (72.3 hr) are shown for comparisons. The landscapes at different ages have been vertically shifted for a better visualization. The color bar shows the relative quasi-potentials in the contour maps. A relative color scheme (blue for low and red for high) is applied in landscapes.

We further examined the PIPL on both the node degrees and normalized RLSs (Figure 2b). The interactions between hub and essential proteins of the old cells at 72.3 hr fell into a deep basin with low relative quasi-potential (ΔU = –94). However, those at younger age from 10.7 to 26.8 hr were located in ridges. Moreover, the hub-versus-essential interactions in the PIPL at 22.0 hr were on a relatively high ridge of ΔU = 60. This analysis indicated that the hub proteins and essential proteins, though they are statistical correlated, may play different role in aging.

### PIPL on dimensions of RLS versus protein turnover

The inversed trends between using the protein abundances and transcript levels may be associated with the protein aggregation during the aging process^27,28^. It may also be owing to the fact that many proteins in yeast have long half-lives^29^. A significant amount of proteins underwent non-exponential degradations in the cells and that the protein degradation rate seemed to be age-dependent^30^. The protein turnover rates were correlated protein functions and activities^31^, and the long-lived proteins may accumulate to high abundances in old cells even though their expression levels had been declined. Here, two independent protein turnover data sets, the CellSys2017 set^31^ and the CellRep2014 set^29^ had been used to construct the age-dependent PIPLs on the dimensions of normalized RLSs versus protein turnover rates (half-lives). Considerable differences can be found between the results of both sets (Figure 3). However, they both interpreted that the short-turnover proteins, which may exhibit higher activities than the long-turnover proteins^31^, had larger fluctuations on the PIPLs of aging cells, especially in interactions with essential proteins. Note that log10-scales had been applied for the turnover rates, in consistent with their distributions^29^.

**Figure 3.**
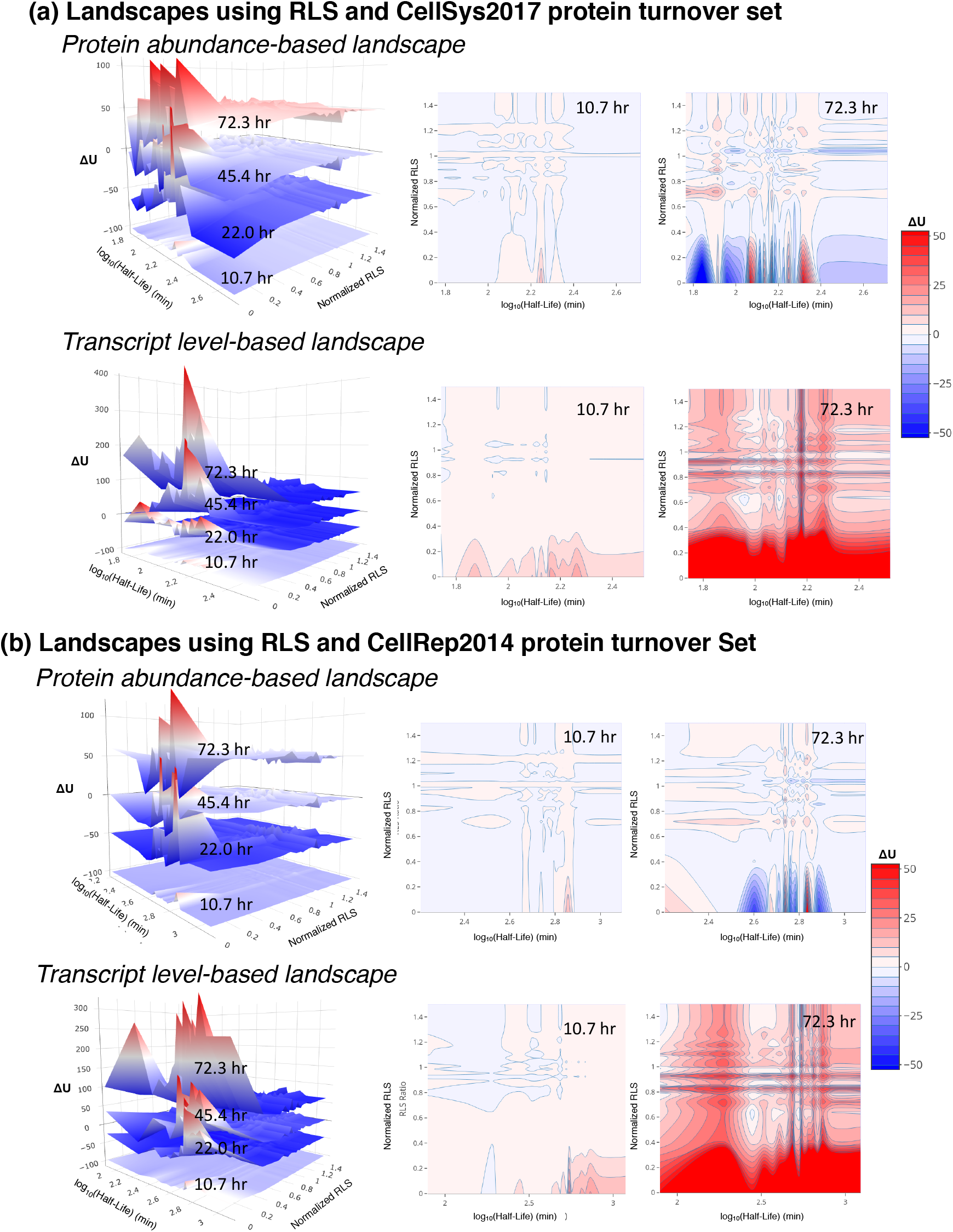
Interactions between essential proteins and short-lived proteins are noisy on the age-dependent PIPLs on dimensions of normalized RLSs and protein turnover rates. The protein turnover data were taken from (**a**) the CellSys2017 set and (**b**) the CellRep2014 set, respectively. The landscapes from both sets showed significant differences between the protein abundance-based (top panels) and transcript level-based (bottom panels) PIPLs. The contour maps of the landscapes of the aging cells at young (10.7 hr) and old (72.3 hr) are shown for comparisons. The landscapes at different ages have been vertically shifted for a better visualization. The color bar shows the relative quasi potentials in the contour maps. A relative color scheme (blue for low and red for high) is applied in landscapes.

Based on the transcript levels (bottom panels), the landscapes using both turnover sets showed elevated quasi-potentials, especially at old age (Figure 3). However, based on protein abundances (top panels), fluctuations of both elevations and declines in the quasi-potentials had been found, consistent with the uncoupling trend between protein and mRNA levels^22^. Nevertheless, in all landscapes we observed large fluctuations attributed to the essential genes (normalized RLS of 0) and their interactions with the low-turnover proteins. The interactions between essential genes and high-turnover proteins are relatively stable during aging, according to the protein-abundance-based PIPLs, but not in the gene-expression-based PIPLs.

Using the above landscapes, we noted that aging has a higher impact to the transcripts (mRNA) levels than to the protein abundances, with respect to the protein-protein interactions. For example, whereas based on the transcript levels for both turnover sets, the interactions with essential genes from all gene groups with different turnovers were in high potential, low probability states, in the protein abundance-based landscapes the interactions between certain groups of genes and the essential genes were dropped to low potential, high probability basins. Moreover, using the CellSys2017 set the low-turnover proteins also showed dropped quasi-potentials or increased interaction probabilities with the essential genes.

### A proteome-wide protein interaction probability landscape based on network permutations

To further understand why lower relative quasi-potential (or higher probability) have been found in the essential-versus-essential interactions based on the proteomics, despite the transcript level trend predicted the opposite direction, we constructed another proteome-wide, global landscape model which calculates the probabilities of that the empirical PIN having more interactions than the random null models. As for the age-dependent landscape models, quasi-potentials were calculated as the negative logarithm of the probabilities to infer basins and hills on this landscape. Similar method had been previously applied to a smaller yeast PIN and observed that the high-degree, hub proteins tend to avoid interactions with the other hub proteins, which was suggested to contribute to the stability of the PIN^32^. This approach had been recently applied to estimate the association strengths between different categories of proteins using a human PIN^16^. Figure 4 shows the global landscape in the variable spaces of node degrees and RLS ratios (i.e., normalized RLS) of single-gene deletion mutations.

**Figure 4.**
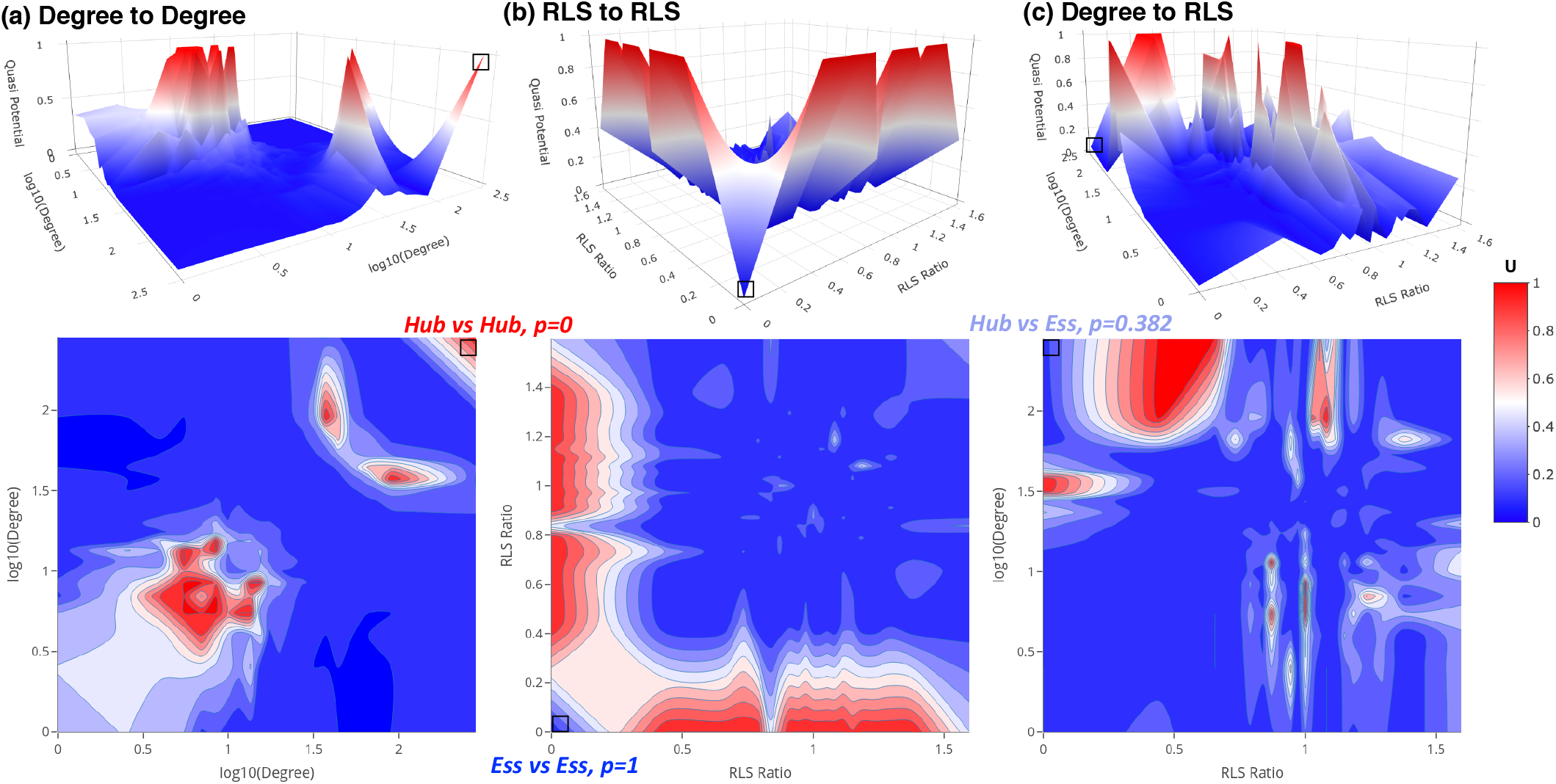
The proteome-wide protein interaction probability landscapes reveal differences between hub and essential proteins. (**a**) The protein interaction landscape on space of the node degrees showed that the hub proteins avoid interactions with other hub proteins (*p* = 0). (**b**) The essential proteins favor interactions with other essential proteins (*p* = 1). (**c**) A moderate ratio of hub-essential interactions (*p* = 0.382) was observed. The hub-hub, essential-essential and hub-essential interaction regions are highlighted as squares on the landscapes and contour maps. The probabilities *p* indicates the ratio of null network models that showed less interactions than the empirical PIN: *p* = 1 (such as the essential-essential interactions) indicates the PIN has more interactions than all null models and *p* = 0 (such as the hub-hub interactions) indicates the PIN has less interactions than all null models, respectively. The quasi potential is calculated as U = −log*p*, the base of the logarithm function is chosen such that U is in range of [0,1]. 22,188 null network models have been used in this work and the *p* = 0 (i.e., *p* < 1/22,188) ratios were converted to *p* = 1/22,189 for estimation of the quasi potential, U. Top panels show the landscapes and bottom panels show the 2D contour maps. The color bar for both the landscapes and contours are shown at right.

The network permutation-based landscape on the node-degree space showed that the hub proteins (high-degree proteins in the PIN) tend to avoid interactions with other hub proteins yet favor the interactions with non-hub proteins (Figure 4a), in agree with previous reports from a smaller yeast PIN^32^. Originally observed in yeast PIN, it had been proposed that the hub-proteins in the PIN tend to be essential proteins and vice versa^26^. This centrality-lethality rule (or hypothesis) had been confirmed in the PINs of other organisms^33^ as well as the genetic interaction network (GIN) of yeast^34^. On the global landscape on the RLS space, however, the global landscape indicated that the essential proteins strongly favor the interactions with other essential proteins but tend to avoid the interactions with non-essential proteins (Figure 4b). The probability of the interactions between essential proteins and hub proteins were moderate (Figure 4c). Therefore, the essential proteins and hub proteins should not be treated equally in these landscapes. The above observation might be conflict with the well-known centrality-lethality rule^26,33,35^.

It is worth noting that the essential genes did have higher degrees (average of 63.3) than non-essential genes (average of 32.7). Based on the landscape on the node degree-RLS space (Figure 4c), we also observed that the interaction densities between hub proteins and essential proteins are significantly stronger than those between the hub proteins and nonessential proteins, also as shown in Figure S1 in the supplementary materials. However, despite of a higher degree in the PIN, a hub protein is not necessarily essential. For example, the most-connected hub protein *DHH1* (YDL160C) has 3,605 interactions in the PIN but is nonessential and the Δ*DHH1* mutant has a normalized RLS of 0.825 compared to the wild type.

### Essential hub genes have increased protein abundances in aging cells

The yeast genome has 5,191 verified, 725 uncharacterized and 688 dubious ORFs (as of 9/17/2020) on the *Saccharomyces* Genome Database^36^. These ORFs encode ca. 1000 essential genes, with any one of which deleted the cell will not be viable^37–40^. We compared the abundances of 4 categories of proteins (genes) including essential hub, essential non-hub, nonessential hub and nonessential non-hub proteins. The protein or transcript abundances were referenced to the ground state, the young cells with age of 7.8 hr, consistent with the PIPLs shown above. We used the node degree k ≥ 100 as a cutoff (which covered the top 5.8% high connectivity proteins) to classify the hub and non-hub proteins (Figure 5a). The yeast life cycle length of 100 minutes was used to estimate the RLS^41^.

**Figure 5.**
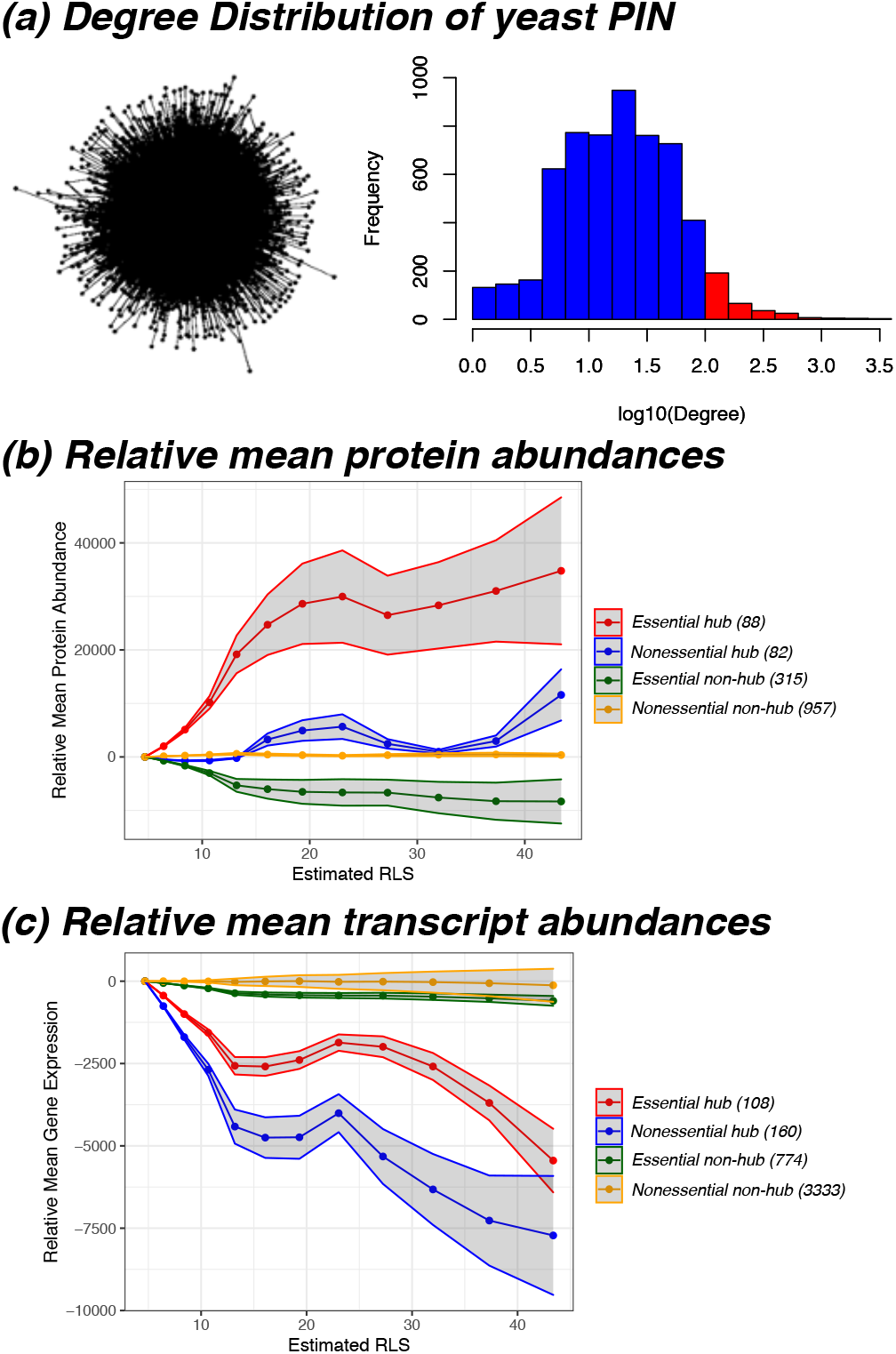
Essential hub proteins (or genes) may account for the opposite trends on the aging landscapes from proteomes and transcriptomes. All proteins (genes) were put in four categories including essential hub (red), nonessential hub (blue), essential non-hub (yellow) and nonessential non-hub proteins or genes. (**a**). Distribution of the node degrees in the yeast PIN. A snapshot of the PIN was shown at left. The hub proteins (red bars) were selected as the node degree k ≥ 100, which were the top 5.8% high-connectivity proteins of the PIN. (**b**). Relative mean protein abundances. (**c**). Relative mean transcript abundances. Average values relative to the young cells at 7.8 hr had been used. The cell ages were approximated to RLS using the dividing time of 100 minutes. The shade of each curve was calculated as the standard deviation from all proteins (genes) of the ratios of aging cells to the ground state, multiplied by the mean differences. The number of proteins (genes) used in the curves of each category had been listed.

Consistent with the evaluation of the relative-quasi potentials (Methods and Materials), the node degrees were weighted to the protein or transcript abundance values in Figure 5b and 5c. We observed that, at old age, the abundances of essential hub proteins increased considerably in the aging cells; however, the abundances of the essential non-hub proteins decreased (Figure 5b). The opposite trends of the average essential protein abundances versus essential transcript abundances (Figure 1) can be explained by the differences in the PIPL related to the essential proteins or transcripts. The differences between the average hub proteins and hub transcripts (Figure 2) can also be explained here. We also listed the numbers of proteins (Figure 5b) or transcripts (Figure 5c) that were included in both the PIN the aging proteomics and transcriptomics data set. These trends of protein and/or transcript abundances may partly explain the basins and ridges observed in the protein interaction probability landscapes. Note that the absolute ages of the cells had been approximated to RLSs using an average division time of 100 minutes. This conversion may not be accurate, for example, the actual division time may increase during aging^42^, and the long estimated RLS in Figure 5 (>40) made it possible that some aging cells may have stopped division and stayed in the post-diauxic, quiescent phase^2^.

### Decline of the PPI dissociation constant may alter the landscapes

All PIPLs shown above were constructed using constant protein-protein interaction dissociation constants (*K*), i.e., the decay rate (d) is 0 for all PPIs. However, in reality, *K* might be affected or altered by many factors, such as post-translational modifications, pH, ionic strengths and other cofactors. It is difficult to measure the variations of *K* for each of the protein-protein interactions. We used an exponential decay model aiming to estimate the potential topological changes of the landscapes attributed to the variations of the dissociation constants, *K*.

As shown in Figure 6, in old yeast cells, the intra-essential interactions arose to a ridge (instead of falling to a basin) when *K* declines with a decay rate of 10^−3^—at this rate *K*_*t*_ becomes half of *K*_*0*_ after 300 cell cycles. We noticed that the dissociation constant strongly affects the essential-essential interactions, partly contributed by their dense interactions. For example, in the proteome-based PIPL, the number of essential-essential interactions (n in equation 7) is 3,465, ~10 times of that between essential proteins and any of the other 20 protein sets based on the quantiles of the normalized RLS.

**Figure 6.**
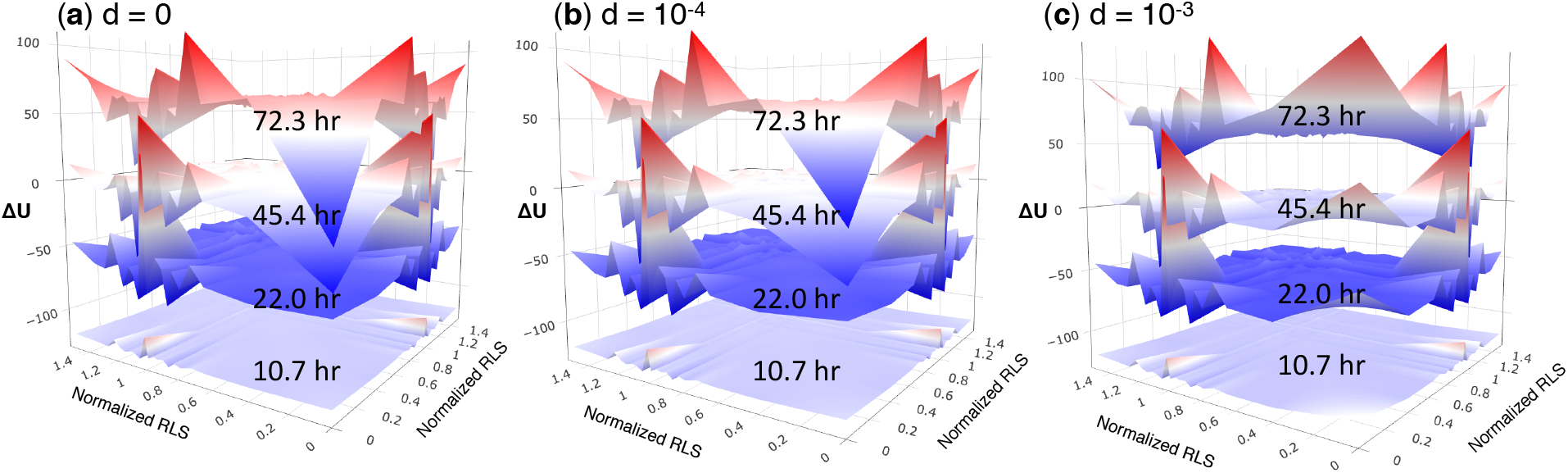
Proteome-based PIPL on RLS with dissociation constants exponentially decay. The decay rate d (in unit of 1/t) of the protein-protein interactions (PPIs) was chosen as (a) d = 0, from which *K*_*43*_ = *K*_*0*_, also see Figure 1; (b) d = 10^−4^, from which *K*_*43*_ = 0.990 *K*_*0*_; and (c) d = 10^−3^, from which *K*_*43*_ = 0.906 *K*_*0*_, where the dissociation constants *K*_*43*_ (at the 43^th^ generation or ca. 71.7 hr, close to the oldest cells in the analysis) was compared with the initial *K*_*0*_. The model with d = 0 (a) corresponds to constant dissociation constants in the models, which was applied to other PIPLs in the main text. However, there is a transition from d = 10^−4^ (b) to d = 10^−3^ (c), in the latter PIPL the intra-essential interactions were located in a ridge for the old cells whereas in the former the intra-essential interactions were also in a basin for the old cells.

The protein interaction probability landscapes on space of other parameters can also be constructed using our approach. As an example, the morphology parameters from the SCMD2 database^38^ had been used in the PIPL as shown in Figure S2 of the supplementary materials.

## Discussion

The role of gene/protein networks in aging have been discussed extensively^21,43–48^, and the role of weakest links in aging were recognized^43,45,46^. A recent mathematical model suggested a role of essential genes in aging^21^. Loss of proteostasis is also a hallmark of aging^49^. The present work on the protein interaction potential landscape during aging suggest that essential hub genes and their interactions may be a critical factor of proteostasis and/or aging. Our results here pose an interesting hypothesis that essential hub genes might be one of the weakest links during aging, a question that could be addressed by future experimental studies.

The protein interaction probability landscapes of the budding yeast in present work used the proteomic and transcriptomic data of aging cells mathematically unmixed from bulk cell culture with young daughter cells and dead cells^22^. We understand that potential biases may persist in the data set and hence in our interpretations. However, using these landscapes we showed that the aging dynamics could be visualized to identify potential aging factors, such as the variations caused by the essential hub proteins. Future study using single cell protein and/or transcript abundance data along aging could be combined with our approach to understand the aging landscape. Moreover, the probability-based interaction landscapes can also be applied using other networks—such as the genetic interaction networks—in understanding of the genotype-phenotype relationships^34,50^.

## Methods and Materials

### Protein-protein interaction network

The empirical protein-protein interaction network (PIN) of S. cerevisiae was obtained from a recent version BioGRID v3.5.177, downloaded on Oct. 1, 2019^23^. Only the physical interactions related to proteins (excluding the self-interactions) were selected from this PIN, comprising 110,290 interactions among 5,784 proteins.

### Network permutation-based association study (NetPAS)

The PIN used in present work is regarded as a simple graph, i.e., undirected graph with no self-loops or multi-interactions. We recently proposed the network permutation-based association study (NetPAS) approach to evaluate the interaction strength between different protein (gene) sets by comparing the original PIN and its random null models^16^. 22,188 null network models were built in a way such that all vertices were randomly reshuffled, yet the node degrees preserved as the original PIN. These network models were also termed the MS02 null models^16,32,51^.

### Replicative lifespan data

The replicative lifespans (RLSs) of 4,698 single-gene deletion strains of *S. cerevisiae* under the YPD (2% glucose) culture had been reported^1^. In addition, the RLSs of single-gene deletion strains under dietary restriction (DR, 0.05% glucose) had been reported^9^. 3,939 genes with RLS data under YPD and 342 genes with RLS data under DR were included in the yeast PIN used in present work, providing the opportunity of constructing the protein interaction landscapes of replicative aging.

### Proteomics and transcriptomics data during aging

One set of proteomics and transcriptomics data of *S. cerevisiae* came from a recent report, which comprises the abundances of 1,494 proteins and expression levels of 4,904 genes collected at 12 different ages ranging from 7.8 to 72.3 hours^22^. We used the young cells at 7.8 hours as the reference (ground state) to evaluate the PIPLs of the older cells at the other 11 ages.

### Protein turnover time

Two sets of in vivo protein turnover time or half-lives have been used. The CellSys2017 set^31^ comprised of 3,160 proteins with 1,341 in the aging proteomics and 2,526 in the aging transcriptomics data sets. The CellRep2014 set^29^ included 3,715 proteins with 1,331 in the aging proteomics and 2,988 in the aging transcriptomics data sets.

### Morphologies of single-gene deletion mutants

We also generated the PIPLs on dimensions of morphological parameters and the RLS of the yeast single-gene deletion mutations. See in the supplementary materials. The morphology data were taken from the SCMD2 (Saccharomyces cerevisiae morphological database 2)^38^. Specifically, two morphologies related to a recent morphological landscape of yeast replicative aging had been analyzed in the protein interaction landscapes^10,52^, including the cell size ratio (C118_C) and bud axis ratio (C114_C) at the late stages. Although the definition of the axis ratio in SCMD2 was different than that used in the morphological landscape, it could provide a rough identification of the roundness of the cell shapes.

### Essential genes of S. cerevisiae

The essential gene set of *S. cerevisiae* were taken from the SCMD2^38^ that comprises 1,112 genes, 1,033 of which have interactions recorded in the PIN.

### Selection of gene groups

We used quantiles to catalog the gene groups in the PIPLs. The yeast PIN covered 3,939 genes with the RLS records of the single-gene deletion mutations. The normalized RLS refers to the RLS ratios between the mutant and the control wide type strain and had been adopted in the landscapes. Among the genes covered by the PIN, 926 non-essential and 403 essential genes had the age-dependent protein abundance records, and 2,991 non-essential and 882 essential genes had the age-dependent transcript records, respectively. In both the age-dependent protein abundance and transcript level sets, the non-essential genes were evenly classified into 20 groups based on the quantiles of the normalized RLSs, and the essential genes were classified into one group with the normalized RLS of 0. For the protein node degrees, half-lives and morphology data sets, all genes were classified into 20 evenly distributed quantiles based on the respective measures. The same classification approach for the RLS groups were applied to the global protein interaction landscape, and all genes included in the PIN and with normalized RLS data had been used.

### Quasi-potentials of age-dependent protein interaction probability landscape

Aging is a dynamic process reflected in the variations of both protein abundances and transcript levels along the aging process. It had been suggested that a configuration of the cell state could formalized as a vector

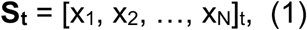

Where N is the total number of genes and xi is the expression level of the i-th gene^53^— we suggest xi could also be the protein abundance of the i-th protein. Under this definition, the aging process could be regarded as the vector variations along time.

For a simple organism that has two interacting proteins x and y:

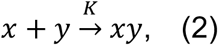

the concentration of the complex xy at time t can be calculated as

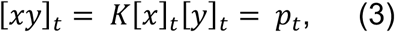

where *K* is the dissociation constant and [x]_t_ and [y]_t_ is the concentration of free x and y at time t; i.e., the probability of forming the complex xy at time t, *p*_*t*_, is approximated as the concentration of [xy] in the cell via equation (3) according to the law of mass action.

The variations of *K* during aging could be described by a simple exponential model

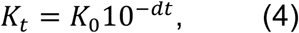

in which *d* is the decay rate, and t is time (age), and *K*_t_ and *K*_0_ are the dissociation constant at age t and 0, respectively (where 0 could be selected as the ground state). The decay rate may be affected by posttranslational modifications, pH and ionic strength variations during aging, and other factors. The dissociation constant *K* remains unchanged during aging if d is 0; *K* decays exponentially if d > 0 and increases exponentially if d < 0. Under this condition and by using the young cell at time 0 as the reference state or ground state, we could get:

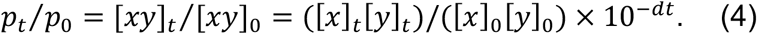

We used a quasi-potential, the negative logarithm of the probabilities, to describe the barriers that the cells must overcome for the transition from one attractor state to the other^53,54^. The quasi-potential^53^ can be calculated from the probabilities using

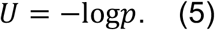

Because all probabilities here are relative to the ground state, the relative quasi potential U_t_ using U_0_ as a reference can be calculated as

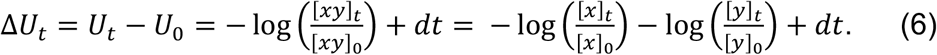

Equation (6) provides a convenient way to estimate the relative quasi potentials at time t, compared to the young cell ground state.

In this work, the relative quasi potential of interactions between two categories of proteins (or genes) are evaluated. The probability is not additive as we cannot sum up the probabilities of different interactions. However, the logarithm function and hence the quasi potential U is additive. Therefore, it is straightforward to calculate the relative quasi potential of interactions between two sets of genes X and Y based on equation (6). For instance, assuming that there are n interactions between X and Y recorded as (x_i_, y_i_), *i* = 1, 2, ..., the relative quasi-potential from all interactions between X and Y would be

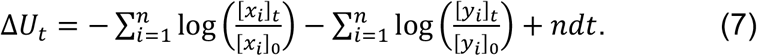

The base of the logarithm function (equations 5–7) is 10 in present work. It should be noted that in present work all quasi-potentials of every single interaction were relative to the ground state and could therefore bear any real values. The unit of the protein abundances or mRNA levels (such as FPKM) were canceled out and did not affect the relative quasi-potentials in equations (4)–(7). In the landscapes of this work, we observed high relative quasi-potentials especially when the interactions between two categories of proteins were dense. However, a cutoff of ±50 was applied in the quasi-potentials of all contour maps because it could capture the indispensable information yet provides relatively good comparison among different landscapes. The variations of the dissociation constant cannot be justified in this work and in above landscapes d was set as 0 (i.e., *K* does not change). However, we showed that significant changes of the landscape with a slow decay rate of the dissociation constant (Figure S3 of the supplementary materials).

### Quasi-potentials of proteome-wide protein-interaction probability landscape

Here we calculated the protein-interaction quasi potentials based on the probability (*p*) for more interactions between certain categories of genes in the PIN than in the null models. The NetPAS method^16^ was used to calculate the *p*-values using 22,188 null models. The protein-interaction quasi potential can also be calculated using equation 5. The base of the logarithm function may vary for the conceptual barriers in equation 5. As 22,188 null models were used, the occurring *p* = 0 cases could be interpreted as *p* < 1/22,188, which was converted to *p* = 1/22,189 in practice to avoid the infinite log0 values. The resulted quasi-potentials were then normalized to [0,1] for the landscape.

## Acknowledgments

This work is supported by NSF Career award #1453078 (transferred to #1720215), BD Spoke #1761839, and internal funding of the University of Tennessee at Chattanooga. All simulations are performed using the SimCenter computing resources of the University of Tennessee at Chattanooga. We thank Dr. Matt Kaeberlein for sharing the replicative lifespan data set of the yeast single-gene mutations with us.

## Supplementary Materials

**Figure S1.**
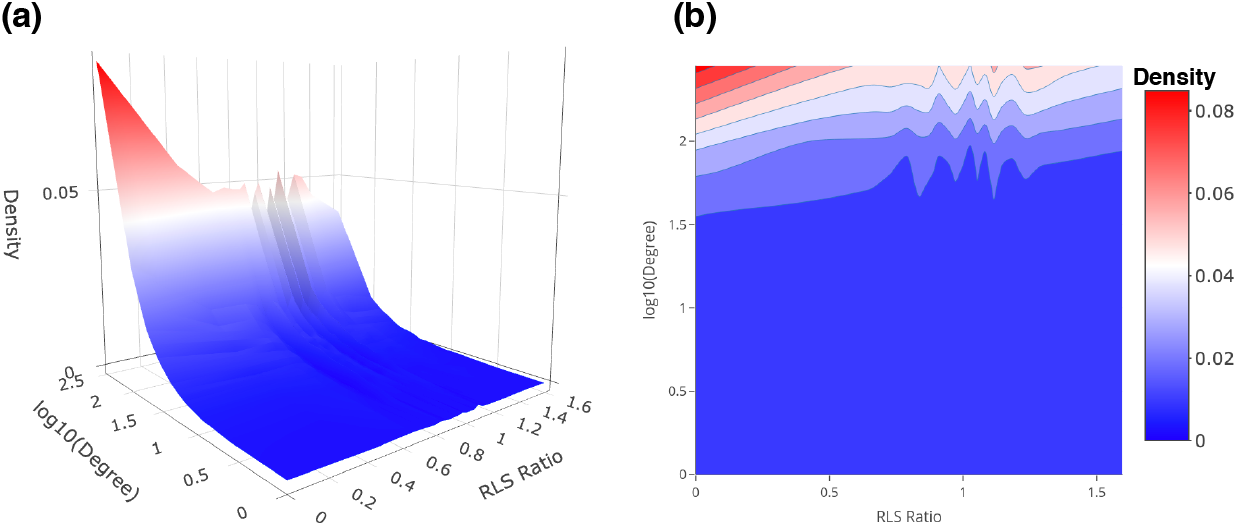
High interaction density was found between essential and hub proteins. The average interaction density calculated on dimensions of node degree and RLS ratios (normalized RLSs) are shown in (**a**) landscape and (**b**) contour map. The interaction density between two sets of nodes is defined as the total number of interactions divided by the by the product of the sizes of both sets.

**Figure S2.**
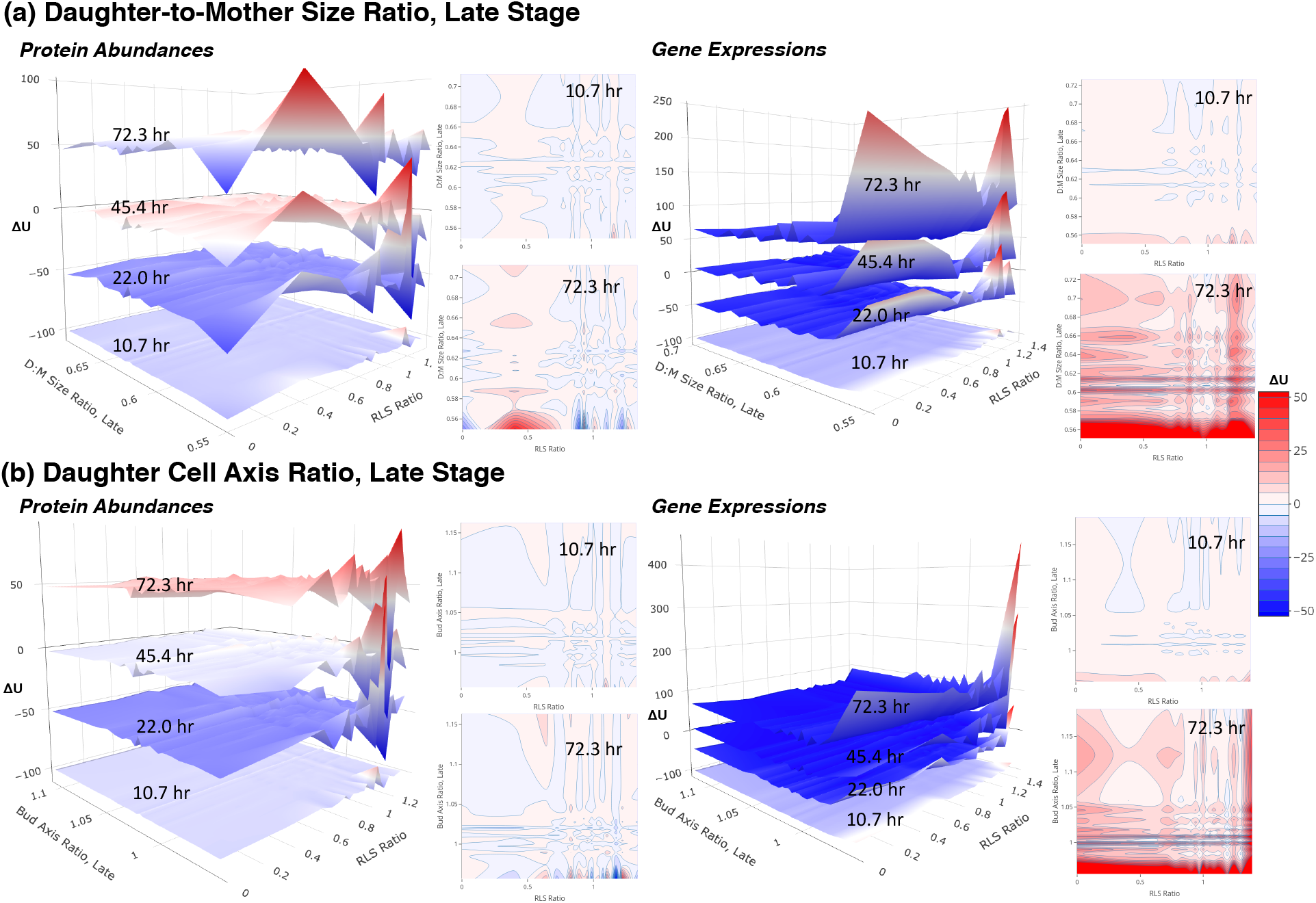
The PIPLs on dimensions of RLS and morphologies. The PIPLs were based on protein abundances (left) or gene expressions (right). The morphologies include (**a**) The daughter-to-mother cell size ratios, and (**b**) the daughter-to-mother cell size ratios. Both morphologies were measured at the late stage of the daughter cell growth, details could be found in the SCMD2^38^. The landscapes at different ages have been vertically shifted for a better visualization. The color bar shows the relative quasi-potentials in the contour maps. A relative color scheme (blue for low and red for high) is applied in landscapes.

A recent work proposed a morphological landscape of yeast aging on which the mother cells took two distinguishable aging paths: in the first path the mother cells produced enlarged and elongated daughter cells, and in the second path the mother cells produced small and round daughter cells^52^. The mother cells went through the first path had longer RLSs than the mother cells underwent the second path. Two morphologies were used in this landscape to infer the aging paths including the daughter-to-mother size ratios for daughter cell size and the axis ratios of the daughter cells for daughter cell roundness, both at the late stages of cell growths^10,52^. Here, we combined the RLSs and took advantage of the Saccharomyces Cerevisiae Morphological Database (SCMD2) which recorded 501 morphologies of different yeast single-gene deletion mutants^38^. Figure S1 showed both the protein abundance-based and gene expression-based PIPLs of RLS versus morphologies obtained from the SCMD2 including the daughter-to-mother size ratios (C118_C) and daughter axis ratios (C114_C) at the late stages of cell growth from the SCMD, both had been used in the construction of the morphological landscape^52^.

Obvious differences between the landscapes based on protein abundances and those based on gene expressions could be found, similar to the landscapes shown above. It should be noted that the morphologies selected from SCMD2 were based on high resolution images of single-gene deletion mutations^38^ such that were different than those in the recent morphological landscape^10,52^. Moreover, the morphological differences in the early and late stages of the daughter cell growth had also been considered ^38^. We did observed differences of the absolute values of the morphologies of cells at early growth stage versus those at late growth stage, however, the orders of these genes were not necessarily conserved. These landscapes could therefore be useful to interpret the general trends related to the morphologies. For example, at the old age, the genes related to small daughter cells (Figure S1) seem to have larger fluctuations in the interactions with other proteins.

The landscapes from present work described the protein-interaction probabilities (or relative quasi-potentials) of single-gene deletion mutants, and they did not represent cell states. Therefore, these landscapes did not predict how the morphological variations affected the RLS of mother cells as the cell-state landscapes^10,52^. Instead, the landscapes of present work suggest that the PINs in the old cells had been altered by aging. To rejuvenate to the young cell state, the old yeast cells needed to redistribute the interactions in the PIN, especially from essential genes, to flatten the above age-dependent landscapes.

## References

1 McCormick, M. A. et al. A comprehensive analysis of replicative lifespan in 4,698 single-gene deletion strains uncovers conserved mechanisms of aging. Cell metabolism 22, 895–906 (2015).

2 Longo, V. D., Shadel, G. S., Kaeberlein, M. & Kennedy, B. Replicative and Chronological Aging in Saccharomyces cerevisiae. Cell Metabolism 16, 18–31, doi:10.1016/j.cmet.2012.06.002 (2012).

3 Janssens, G. E. & Veenhoff, L. M. Evidence for the hallmarks of human aging in replicatively aging yeast. Microb Cell 3, 263–274, doi:10.15698/mic2016.07.510 (2016).

4 Crane, M. M. & Kaeberlein, M. The paths of mortality: how understanding the biology of aging can help explain systems behavior of single cells. Current opinion in systems biology 8, 25–31 (2018).

5 Wasko, B. M. & Kaeberlein, M. Yeast replicative aging: a paradigm for defining conserved longevity interventions. Fems Yeast Res 14, 148–159, doi:10.1111/1567-1364.12104 (2014).

6 Defossez, P. A. et al. Elimination of replication block protein Fob1 extends the life span of yeast mother cells. Mol Cell 3, 447–455, doi:10.1016/s1097-2765(00)80472-4 (1999).

7 Guo, Z. H., Adomas, A. B., Jackson, E. D., Qin, H. & Townsend, J. P. SIR2 and other genes are abundantly expressed in long-lived natural segregants for replicative aging of the budding yeast Saccharomyces cerevisiae. Fems Yeast Res 11, 345–355, doi:10.1111/j.1567-1364.2011.00723.x (2011).

8 Kaeberlein, M. & Powers, R. W. Sir2 and calorie restriction in yeast: A skeptical perspective. Ageing Res Rev 6, 128–140, doi:10.1016/j.arr.2007.04.001 (2007).

9 Schleit, J. et al. Molecular mechanisms underlying genotype-dependent responses to dietary restriction. Aging Cell 12, 1050–1061, doi:10.1111/acel.12130 (2013).

10 Li, Y. et al. A programmable fate decision landscape underlies single-cell aging in yeast. Science 369, 325–329, doi:10.1126/science.aax9552 (2020).

11 Sen, P. et al. H3K36 methylation promotes longevity by enhancing transcriptional fidelity. Genes Dev 29, 1362–1376, doi:10.1101/gad.263707.115 (2015).

12 Ke, Z. et al. Translation fidelity coevolves with longevity. Aging Cell 16, 988–993, doi:10.1111/acel.12628 (2017).

13 Lahtvee, P. J. et al. Absolute Quantification of Protein and mRNA Abundances Demonstrate Variability in Gene-Specific Translation Efficiency in Yeast. Cell Systems 4, 495–+, doi:10.1016/j.cels.2017.03.003 (2017).

14 Hartwell, L. H., Hopfield, J. J., Leibler, S. & Murray, A. W. From molecular to modular cell biology. Nature 402, C47–C52, doi:Doi 10.1038/35011540 (1999).

15 Yu, H. Y. et al. High-quality binary protein interaction map of the yeast interactome network. Science 322, 104–110, doi:10.1126/science.1158684 (2008).

16 Guo, H. B. & Qin, H. Association study based on topological constraints of protein-protein interaction networks. Sci Rep 10, 10797, doi:10.1038/s41598-020-67875-w (2020).

17 Barabasi, A.-L. & Oltvai, Z. N. Network biology: understanding the cell’s functional organization. Nature reviews genetics 5, 101 (2004).

18 Gavin, A. C. et al. Proteome survey reveals modularity of the yeast cell machinery. Nature 440, 631–636, doi:DOI 10.1038/nature04532 (2006).

19 Xue, H. L. et al. A modular network model of aging. Molecular Systems Biology 3, doi:10.1038/msb4100189 (2007).

20 Faisal, F. E. & Milenkovic, T. Dynamic networks reveal key players in aging. Bioinformatics 30, 1721–1729, doi:10.1093/bioinformatics/btu089 (2014).

21 Qin, H. Estimating network changes from lifespan measurements using a parsimonious gene network model of cellular aging. Bmc Bioinformatics 20, doi:10.1186/s12859-019-3177-7 (2019).

22 Janssens, G. E. et al. Protein biogenesis machinery is a driver of replicative aging in yeast. Elife 4, e08527, doi:10.7554/eLife.08527 (2015).

23 Oughtred, R. et al. The BioGRID interaction database: 2019 update. Nucleic Acids Research 47, D529–D541, doi:10.1093/nar/gky1079 (2019).

24 Ying, Q. L. et al. The ground state of embryonic stem cell self-renewal. Nature 453, 519–U515, doi:10.1038/nature06968 (2008).

25 Chandler, D. Introduction to modern statistical mechanics. (Oxford University Press, 1987).

26 Jeong, H., Mason, S. P., Barabasi, A. L. & Oltvai, Z. N. Lethality and centrality in protein networks. Nature 411, 41–42, doi:Doi 10.1038/35075138 (2001).

27 David, D. C. et al. Widespread Protein Aggregation as an Inherent Part of Aging in C. elegans. Plos Biology 8, doi:10.1371/journal.pbio.1000450 (2010).

28 Saarikangas, J. & Barral, Y. Protein aggregates are associated with replicative aging without compromising protein quality control. Elife 4, doi:10.7554/eLife.06197 (2015).

29 Christiano, R., Nagaraj, N., Frohlich, F. & Walther, T. C. Global Proteome Turnover Analyses of the Yeasts S-cerevisiae and S-pombe. Cell Reports 9, 1959–1965, doi:10.1016/j.celrep.2014.10.065 (2014).

30 McShane, E. et al. Kinetic Analysis of Protein Stability Reveals Age-Dependent Degradation. Cell 167, 803–815 e821, doi:10.1016/j.cell.2016.09.015 (2016).

31 Martin-Perez, M. & Villen, J. Determinants and Regulation of Protein Turnover in Yeast. Cell Systems 5, 283–+, doi:10.1016/j.cels.2017.08.008 (2017).

32 Maslov, S. & Sneppen, K. Specificity and stability in topology of protein networks. Science 296, 910–913 (2002).

33 Raman, K., Damaraju, N. & Joshi, G. K. The organisational structure of protein networks: revisiting the centrality-lethality hypothesis. Syst Synth Biol 8, 73–81, doi:10.1007/s11693-013-9123-5 (2014).

34 Costanzo, M. et al. A global genetic interaction network maps a wiring diagram of cellular function. Science 353, aaf1420 (2016).

35 Peng, X. Q., Wang, J. X., Wang, J., Wu, F. X. & Pan, Y. Rechecking the Centrality-Lethality Rule in the Scope of Protein Subcellular Localization Interaction Networks. Plos One 10, doi:10.1371/journal.pone.0130743 (2015).

36 Cherry, J. M. et al. Saccharomyces Genome Database: the genomics resource of budding yeast. Nucleic Acids Res 40, D700–705, doi:10.1093/nar/gkr1029 (2012).

37 Luo, H., Lin, Y., Gao, F., Zhang, C.-T. & Zhang, R. DEG 10, an update of the database of essential genes that includes both protein-coding genes and noncoding genomic elements. Nucleic acids research 42, D574–D580 (2013).

38 Ohya, Y. et al. High-dimensional and large-scale phenotyping of yeast mutants. Proc Natl Acad Sci U S A 102, 19015–19020, doi:0509436102 [pii] 10.1073/pnas.0509436102 (2005).

39 Rancati, G., Moffat, J., Typas, A. & Pavelka, N. Emerging and evolving concepts in gene essentiality. Nature Reviews Genetics 19, 34 (2018).

40 Dowell, R. D. et al. Genotype to Phenotype: A Complex Problem. Science 328, 469–469, doi:10.1126/science.1189015 (2010).

41 Herskowitz, I. Life-Cycle of the Budding Yeast Saccharomyces-Cerevisiae. Microbiol Rev 52, 536–553, doi:Doi 10.1128/Mmbr.52.4.536-553.1988 (1988).

42 Jo, M. C., Liu, W., Gu, L., Dang, W. W. & Qin, L. D. High-throughput analysis of yeast replicative aging using a microfluidic system. P Natl Acad Sci USA 112, 9364–9369, doi:10.1073/pnas.1510328112 (2015).

43 Skurnick, I. D. & Kemeny, G. Stochastic studies of aging and mortality in multicellular organisms. I. The asymptotic theory. Mech Ageing Dev 7, 65–80, doi:10.1016/0047-6374(78)90053-2 (1978).

44 Kirkwood, T. B. & Kowald, A. Network theory of aging. Exp Gerontol 32, 395–399 (1997).

45 Csermely, P. & Soti, C. Cellular networks and the aging process. Arch Physiol Biochem 112, 60–64, doi:10.1080/13813450600711243 (2006).

46 Soti, C. & Csermely, P. Aging cellular networks: chaperones as major participants. Exp Gerontol 42, 113–119, doi:10.1016/j.exger.2006.05.017 (2007).

47 Promislow, D. E. Protein networks, pleiotropy and the evolution of senescence. Proc Biol Sci 271, 1225–1234, doi:10.1098/rspb.2004.2732 (2004).

48 Farkas, I. J. et al. Network-based tools for the identification of novel drug targets. Science Signaling 4, pt3–pt3 (2011).

49 Lopez-Otin, C., Blasco, M. A., Partridge, L., Serrano, M. & Kroemer, G. The Hallmarks of Aging. Cell 153, 1194–1217, doi:10.1016/j.cell.2013.05.039 (2013).

50 Costanzo, M. et al. Global Genetic Networks and the Genotype-to-Phenotype Relationship. Cell 177, 85–100, doi:10.1016/j.cell.2019.01.033 (2019).

51 Qin, H., Lu, H. H., Wu, W. B. & Li, W.-H. Evolution of the yeast protein interaction network. Proceedings of the National Academy of Sciences 100, 12820–12824 (2003).

52 Jin, M. et al. Divergent Aging of Isogenic Yeast Cells Revealed through Single-Cell Phenotypic Dynamics. Cell Systems 8, 242–253, doi:10.1016/j.cels.2019.02.002 (2019).

53 Huang, S. Reprogramming cell fates: reconciling rarity with robustness. Bioessays 31, 546–560, doi:10.1002/bies.200800189 (2009).

54 Wang, J., Zhang, K., Xu, L. & Wang, E. Quantifying the Waddington landscape and biological paths for development and differentiation. P Natl Acad Sci USA 108, 8257–8262, doi:10.1073/pnas.1017017108 (2011).

